# Inhibitory NK receptors regulate the γδ T cell response to malaria

**DOI:** 10.1101/2025.08.18.670802

**Authors:** Meagan E. Olive, Perri C. Callaway, Mikias Ilala, Justine Levan, Gonzalo R. Acevedo, Felistas Nankya, Emmanuel Arinaitwe, John Rek, Prasanna Jagannathan, Grant Dorsey, Moses R. Kamya, Margaret E. Feeney

**Affiliations:** Department of Medicine, San Francisco General Hospital, University of California, San Francisco, San Francisco, California, United States of America; Biomedical Sciences PhD Program, University of California, San Francisco, San Francisco, California, United States of America; Infectious Diseases and Immunity Graduate Group, University of California, Berkeley, Berkeley, California, United States of America; Infectious Diseases Research Collaboration, Kampala, Uganda; Department of Medicine, Stanford University, Stanford, California, United States of America; Department of Medicine, Makerere University, Kampala, Uganda; Department of Pediatrics, University of California, San Francisco, San Francisco, California, United States of America

**Keywords:** Gamma delta T cells, NK receptors, malaria

## Abstract

Gamma delta (γδ) T cells are important mediators of the immune response to childhood malaria infection. Human Vγ9^+^Vδ2^+^ T cells possess intrinsic, HLA-independent responsiveness to *Plasmodium falciparum* phosphoantigens produced in the blood stage of malaria infection. Engagement of the γδ T cell receptor (TCR) by phosphoantigen-bound butyrophilin molecules results in Vγ9^+^Vδ2^+^ T cell expansion, pro-inflammatory cytokine production, and release of cytotoxic granules that mediate parasite killing. Repeated malaria infection, however, leads to a reduction in circulating Vγ9^+^Vδ2^+^ T cells and upregulation of immunomodulatory molecules, including NK receptors, that correlates with less severe symptoms upon infection. We explore phenotypic and functional differences of γδ T cells in Ugandan children with high versus low malaria exposure, utilizing high-parameter spectral flow cytometry analysis of PBMCs. We observed significant differences in expression of inhibitory NK receptors – KIR2DL1, KIR2DL2/3, KIR3DL1, LILRB1, and NKG2A – on γδ T cell subsets, with Vγ9^+^Vδ2^+^ T cells exhibiting a divergent mechanism of control compared to other subsets. We found that NKG2A and KIR3DL1 expression associated with potent Vγ9^+^Vδ2^+^ T cell responses to TCR- and Fc receptor (FcR)-mediated stimulation while KIR2DL1, KIR2DL2/3 and LILRB1 associated with reduced degranulation and cytokine production. These results identify a new role for inhibitory NK receptors expressed on γδ T cells, exerting a finely tuned balance of activating and inhibitory signals to regulate the response to malaria-related antigens.

**AUTHOR SUMMARY:** Malaria remains one of the deadliest infectious diseases, disproportionately affecting young children in sub-Saharan Africa who succumb to sequelae of *Plasmodium falciparum* infection. However, children living in highly endemic areas experience repeated malaria infection and develop naturally acquired–but not sterilizing–immunity which leads to an asymptomatic reinfection pattern. The immune factors that determine the balance of inflammatory and tolerogenic functions seen in non-sterilizing malaria immunity are yet to be fully understood. Here, we focus on the phenotypic and functional differences in one cell type between children with a history of low versus high malaria exposure. We identified a group of inhibitory surface receptors that improved the antimalarial function of this cell, and another group that worsened their function. Our study clarifies the immune landscape in highly malaria-exposed individuals and illuminates one potential system of regulating the cellular response to repeat infection.

## INTRODUCTION

The Vγ9^+^Vδ2^+^ subset of γδ T cells plays an important role in the immune response to blood-stage malaria. The Vγ9^+^Vδ2^+^ TCR responds to *Plasmodium falciparum* phosphoantigens by recognizing a conformational change of ubiquitous host butyrophilin proteins initiated by phosphoantigen binding—a process that is independent of HLA [1–4]. In acute malaria infection of a naive host, Vγ9^+^Vδ2^+^ T cells expand to comprise up to 15-30% of circulating mononuclear cells, and produce pro-inflammatory cytokines as well as cytotoxic granules containing granulysin, which mediates direct merozoite killing [5–11]. However, the frequency of circulating Vγ9^+^Vδ2^+^ T cells and their pro-inflammatory responsiveness to malaria antigens both decline markedly in individuals following chronic repeated malaria exposure [12,13].

γδ T cells are regarded as “semi-innate”, and in addition to TCR-mediated molecular pattern recognition, they can be rapidly and independently activated via innate mechanisms much like natural killer (NK) cells. These mechanisms include sensing of stress molecules by NKG2D, responding to IL-12/IL-18 via cytokine receptors, and importantly inducing antibody-dependent cellular cytotoxicity (ADCC) upon engagement of the Fc receptor CD16 [14–17]. We have shown that in heavily malaria-exposed individuals, Vγ9^+^Vδ2^+^ T cells frequently upregulate CD16 and become capable of ADCC, despite losing sensitivity to TCR stimulation [18,19]. Rather, CD16-expressing Vγ9^+^Vδ2^+^ T cells respond independently of their TCR to IgG-opsonized *P. falciparum*-infected red blood cells (RBCs) [19]. This transition to a mature NK-like activation program poises Vγ9^+^Vδ2^+^ T cells to maintain effector functions under chronic antigen exposure, enabling a response that utilizes circulating antimalarial antibodies present in later infection [19].

γδ T cells also express other receptors classically associated with NK cells—referred to as NK receptors or NKRs—including inhibitory NKRs such as killer cell immunoglobulin-like receptors (KIR), leukocyte immunoglobulin-like receptor B1 (LILRB1), and NKG2A [20,21]. However, the impact of inhibitory NKR expression on γδ T cell function remains underexplored. In children with heavy malaria exposure, Vγ9^+^Vδ2^+^ T cells that express CD16 more frequently express inhibitory KIR, including KIR2DL1, KIR2DL3, and KIR3DL1 [19]. Further, individuals with HLA-C2 and HLA-Bw4, ligands for KIR2DL1 and KIR3DL1 respectively, have an increased risk of *P. falciparum* parasitemia [22]. The evolutionary importance of these inhibitory signaling pathways is underscored by the fact that malaria targets inhibitory NK receptors for immune evasion. RIFINs (repeated interspersed families of polypeptides), for example, are *P. falciparum* proteins expressed on the surface of infected RBCs that bind and suppress immune cells through inhibitory receptors including LILRB1 and KIR2DL1. Inhibition of NK cells by this interaction correlates with more severe malaria [23,24], likely due to disruption of NK cells’ parasite-killing ADCC function [25]. We hypothesized that inhibitory NK receptors expressed on γδ T cells contribute to the regulation of the response to malaria, dampening the inflammatory response of Vγ9^+^Vδ2^+^ T cells in individuals with chronic repeated infection. Given that our current knowledge of inhibitory NKRs in malaria relies on genetic association data and observations in NK cells, a better understanding of which subpopulations of γδ T cells express inhibitory NK receptors and how this expression affects their response to malaria antigens is necessary.

Here, we used high parameter spectral flow cytometry to examine the expression of inhibitory NK receptors on the major circulating γδ T cell subsets in Ugandan children residing in regions of high and low malaria transmission intensity, and to explore their impact on antimalarial function. We found that Vγ9^+^Vδ2^+^ T cells predominantly express NKG2A, while Vδ1^+^ and Vγ9^−^Vδ2^+^ T cells in peripheral blood more frequently express the inhibitory receptors LILRB1 and/or KIR. In addition, we found that inhibitory NK receptor expression on Vγ9^+^Vδ2^+^ T cells was associated with functional differences in pro-inflammatory cytokine production and degranulation in response to *in vitro* stimulation with malaria-derived phosphoantigens.

## RESULTS

### γδ T cell subset frequencies shift with malaria exposure

To compare γδ T cells from individuals with low versus high malaria exposure, we leveraged samples from children enrolled in an observational cohort study conducted in two regions of Uganda: Jinja (n = 25), a peri-urban district with low malaria transmission, and Tororo (n = 50), a rural district with high malaria transmission [26] (**Fig 1A**). Participants ranged from 4 to 12 years old (mean = 7.8 years) and were balanced by biological sex (Jinja: 52% male, 48% female; Tororo: 60% male, 40% female) (**Fig 1A**). Instances of *P. falciparum* parasitemia, measured by quarterly blood draw in addition to symptomatic malaria visits, were significantly more frequent in children from Tororo (median = 3.8 events per year) compared to Jinja (median = 0 events per year; *p = 1.5e-10*) (**Fig 1A**). The overall frequency of γδ T cells did not differ between the two study sites, constituting approximately 5% of peripheral blood T cells in each group (**Fig 1B, S1 Fig**). However, the distribution of γδ T cell subsets (defined by their γ and δ TCR chain usage) was starkly different, with proportionally more Vγ9^−^Vδ2^−^ T cells and fewer Vγ9^+^Vδ2^+^ T cells in children from Tororo than from Jinja (**Fig 1C**). The Vγ9^−^Vδ2^−^ T cell population likely consists mainly of Vδ1^+^ T cells, the only other major subset observed in peripheral blood beyond the neonatal period [27–29]. Frequencies of the minor γδ T cell subsets (Vγ9^+^Vδ2^−^ and Vγ9^−^Vδ2^+^) did not differ by site. Across both study sites, we observed a direct correlation between the proportion of Vγ9^−^Vδ2^−^ T cells and parasitemic events per year, and a corresponding inverse correlation with the proportion of Vγ9^+^Vδ2^+^ T cells (**Fig 1D**). This pattern is consistent with prior studies showing that in heavily malaria-exposed children, Vγ9^+^Vδ2^+^ T cells are lower in both frequency and absolute number and Vδ1^+^ T cell frequencies are higher [12,13,30].

**Fig 1.**
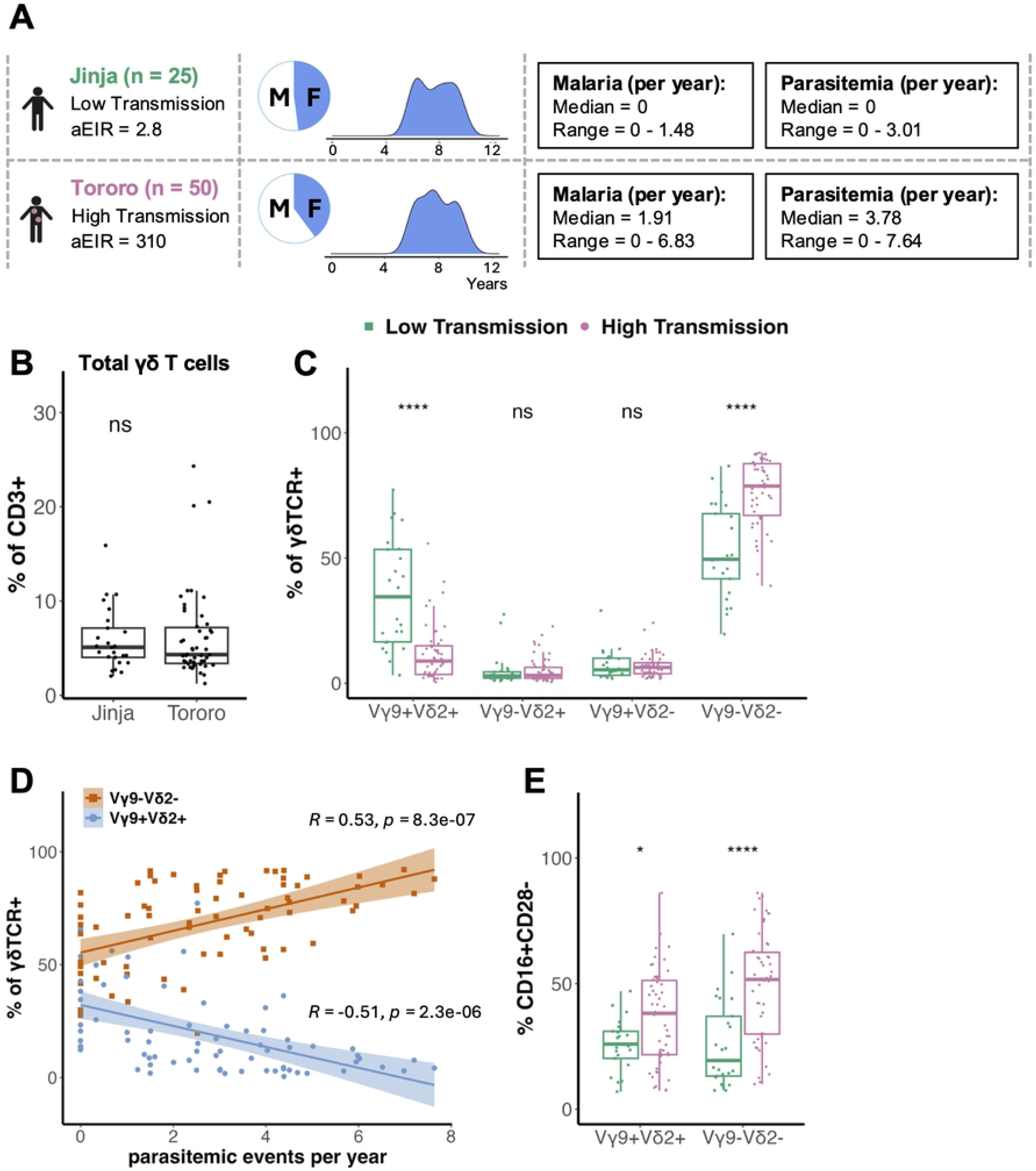
Malaria exposure associates with differences in γδ T cell subset frequency and effector phenotype. (**A**) Participant details from PRISM cohort. Samples were selected from individuals living in two study sites: Jinja (low transmission, n = 25, green) and Tororo (high transmission, n = 50, pink). Annual entomological inoculation rate (aEIR) is at the time of sample collection [26]. Participants had an average age of 7.78 years and were balanced by sex. Malaria episodes and parasitemic events were quantified as a yearly value for each participant and reported as a group median and range. *Ex vivo* frequency of (**B**) total γδ T cells (pan-γδTCR^+^) out of CD3^+^ T cells, (**C**) γδ T cell subsets out of pan-γδTCR^+^ T cells as defined by expression of Vγ9 and/or Vδ2 TCR chains, and (**D**) Vγ9^+^Vδ2^+^ (blue) and Vγ9^−^Vδ2^−^ (orange) T cells as a function of parasitemic events per year. Regression line is a standard linear model with the 95% confidence interval shaded. (**E**) CD16^+^CD28^−^ Vγ9^+^Vδ2^+^ and Vγ9^−^Vδ2^−^ T cells in PBMC from individuals living in low and high malaria transmission sites. Study site groups were compared with an unpaired Wilcoxon rank sum test. For all analyses, a 2-tailed P value < 0.05 was considered significant (*: p <= 0.05, ****: p <= 0.0001).

We have previously observed that in heavily malaria-exposed children, Vδ2^+^ T cells frequently upregulate CD16 and become highly differentiated [18,19]. We therefore assessed the proportion of γδ T cells in each subset that adopt a mature, non-naïve (CD28^−^) cytotoxic (CD16^+^) effector phenotype [31]. We found that both the Vγ9^+^Vδ2^+^ and Vγ9^−^Vδ2^−^ subsets had a higher frequency of CD16^+^CD28^−^ cells in highly malaria-exposed Tororo children than in children from Jinja (**Fig 1E**). This suggests that under high antigen load, peripheral blood γδ T cells shift, increasing the proportion of cells that adopt an NK-like phenotype and can be alternately activated by circulating antibodies. We have previously shown that Vγ9^+^Vδ2^+^ T cells that express CD16 are less responsive to activation by malaria-derived phosphoantigens [19]. Furthermore, in highly malaria exposed children, Vδ2^+^ T cells upregulate numerous inhibitory receptors, including those typical of NK cells [12]. Therefore, we next considered whether co-expression of inhibitory NK receptors may alter the phenotypic and functional properties of γδ T cells in response to malaria.

### Inhibitory NK receptor expression differs between γδ T cell subsets

Prior studies have shown that expression of inhibitory NK receptors such as KIR and the NKG2 family of lectin-like receptors are more highly expressed on γδ T cells compared to conventional αβ T cells; however, differences in expression among γδ T cell subsets have not been well described. We measured surface expression of NKG2A, KIR2DL1, KIR2DL2/3, KIR3DL1, and LILRB1 on γδ T cell subsets. We observed two very distinct patterns. Vγ9^+^Vδ2^+^ T cells, which have been described as more innate-like due to their restricted TCR diversity and rapid recognition of microbial patterns [14,16], frequently expressed NKG2A (median = 66.7%), whereas LILRB1 expression was uncommon (median = 8.6%) and expression of all 3 inhibitory KIR was very rare (median < 6%) (**Fig 2A**). In contrast, Vγ9^−^Vδ2^−^, Vγ9^+^Vδ2^−^, and Vγ9^−^Vδ2^+^ T cells, which are considered adaptive [32,33], frequently expressed LILRB1 (median = 58.6%, 41.5%, 49.4%, respectively), but they expressed NKG2A only very rarely in most donors (median < 7%). Expression of all 3 inhibitory KIR was also much higher on these subsets than on Vγ9^+^Vδ2^+^ T cells, with KIR2DL2/3 being the most frequently expressed, followed by KIR3DL1, then KIR2DL1 (**Fig 2A**). These data suggest a divergent program of inhibitory control of Vγ9^+^Vδ2^+^ T cells compared to the other γδ T cell subsets which favors expression of the non-polymorphic NKG2A receptor.

**Fig 2.**
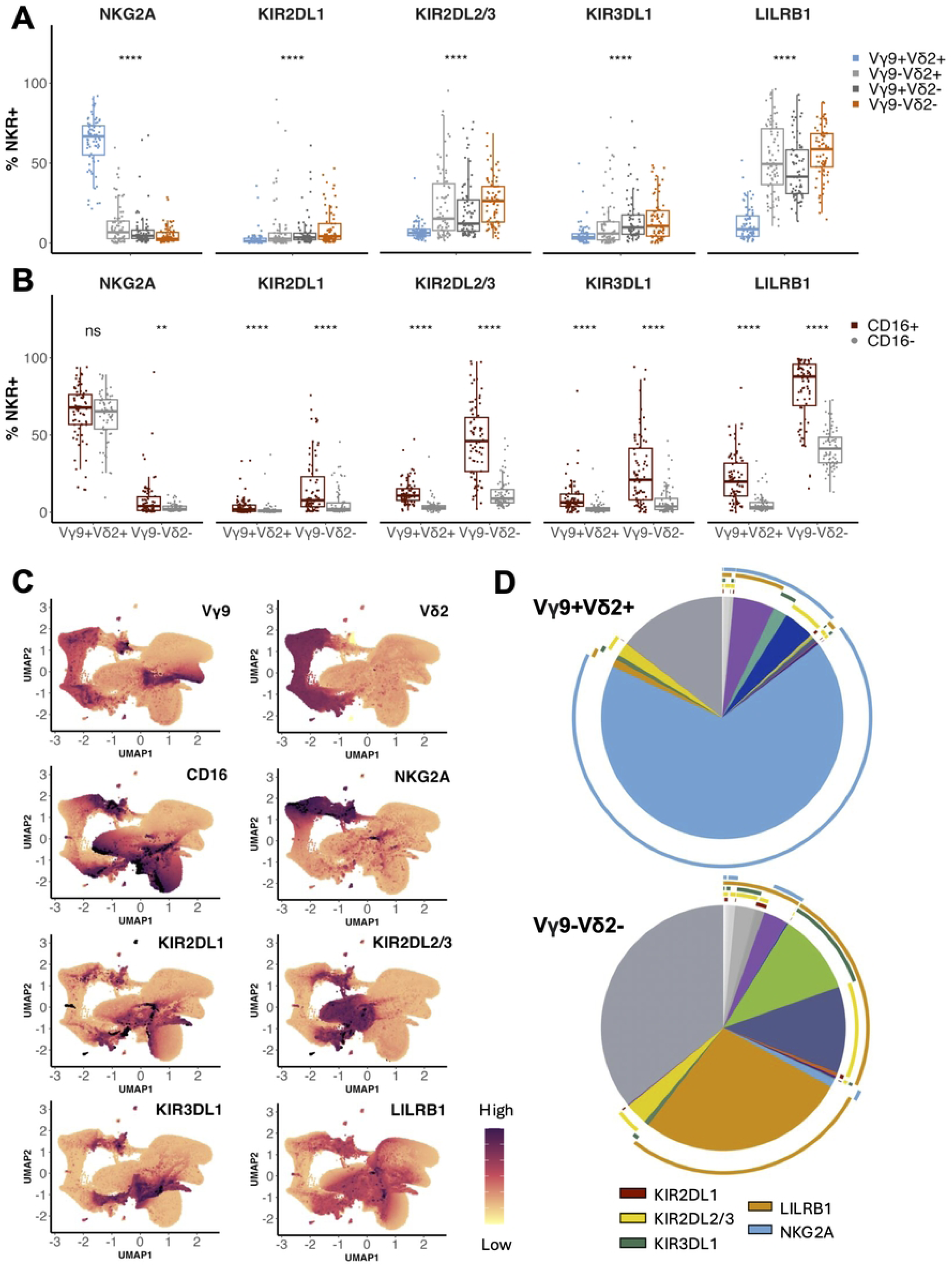
Expression of inhibitory NK receptors differs among γδ T cell subsets. (**A**) *Ex vivo* expression of inhibitory NK receptors—NKG2A, KIR2DL1, KIR2DL2/3, KIR3DL1, and LILRB1—on γδ T cell subsets (Vγ9^+^Vδ2^+^: light blue, Vγ9^−^Vδ2^+^: light gray, Vγ9^+^Vδ2^−^: dark gray, Vγ9^−^Vδ2^−^: orange) in all participants (n = 75). A Kruskal-Wallis test was used to statistically assess the differences in NKR expression across the four γδ T cell subsets. (**B**) Frequency of NKR^+^ cells among either CD16^+^ (maroon) or CD16^−^ (gray) Vγ9^+^Vδ2^+^ and Vγ9^−^Vδ2^−^ T cells. CD16 expression categories were compared with an unpaired Wilcoxon rank sum test. For **A** and **B**, a 2-tailed P value < 0.05 was considered significant (**: p <= 0.01, ****: p <= 0.0001). (**C**) Unsupervised UMAP plots depicting normalized surface marker expression on γδ T cells from all samples. Purple = high expression and yellow = low expression, with the arcsinh-transformed expression ranging from −1 to 5. (**D**) SPICE pie charts illustrating co-expression of NK receptors on either Vγ9^+^Vδ2^+^ or Vγ9^−^Vδ2^−^ T cells. Wedges represent the averaged frequencies of a given phenotypic population across all samples, with colored arcs indicating receptor expression (KIR2DL1: maroon, KIR2DL2/3: yellow, KIR3DL1: green, LILRB1: orange, NKG2A: light blue).

Next, we assessed whether inhibitory NK receptors were preferentially expressed on CD16^+^ γδ T cells, which exhibit NK-like antibody-dependent cytotoxic function [19,34,35]. NKG2A was expressed more frequently on CD16^+^ Vγ9^−^Vδ2^−^ T cells, although its expression overall was low (**Fig 2B**). In contrast, the high expression of NKG2A on Vγ9^+^Vδ2^+^ T cells did not differ between CD16^+^ and CD16^−^ cells (**Fig 2B**). We found the expression of LILRB1 and all three KIR to be much more frequent on CD16^+^ cells compared to CD16^−^ cells in both the Vγ9^+^Vδ2^+^ and Vγ9^−^Vδ2^−^ T subsets (**Fig 2B**).

To determine the extent to which inhibitory NK receptors were co-expressed on individual γδ T cells, we performed high-dimensional analysis [36] and SPICE co-expression analysis [37] of our flow cytometry data. Interestingly, dimensionality reduction and UMAP analysis showed that overall, NK receptor expression was heterogeneous and widely distributed across the population of γδ T cells, rather than overlapping in a distinct NK-like cluster (**Fig 2C**). For example, KIR2DL2/3 and KIR3DL1 expression segregated to different regions of the UMAP while LILRB1 expression was diffuse across most cells. SPICE analysis showed that a sizeable fraction of the Vγ9^−^Vδ2^−^ T cell population co-expressed LILRB1 along with one or more KIR (especially KIR2DL2/3 and KIR3DL1), while co-expression was less common in Vγ9^+^Vδ2^+^ T cells, most of which expressed NKG2A alone (**Fig 2D**). Together, these data suggest that the innate-like Vγ9^+^Vδ2^+^ T cells are mainly regulated by NKG2A regardless of CD16 expression, and the adaptive Vγ9^−^Vδ2^−^ T cells are mainly regulated by LILRB1 in their more mature CD16-expressing state.

### Malaria exposure intensity influences expression of LILRB1, but not KIR or NKG2A

Next, we aimed to determine whether chronic repeated malaria exposure impacts NKR expression. To do so, we first compared NK receptor expression on total γδ T cells between participants residing in the low (Jinja) versus high (Tororo) malaria transmission intensity sites. Overall, the proportion of γδ T cells expressing NKG2A was lower in children with high malaria exposure, while the proportion of γδ T cells expressing LILRB1 was higher, and KIR expression did not differ significantly (**Fig 3A**). However, when assessed on Vγ9^+^Vδ2^+^ and Vγ9^−^Vδ2^−^ T cell subsets, there was no difference in NKG2A or KIR expression between study sites (**Fig 3B**). This indicates that although malaria exposure has a dramatic influence on the relative proportion of γδ T cell subsets in peripheral blood (**Fig 1C**), it does not change the proportion of NKG2A- or KIR-expressing cells within these subsets.

**Fig 3.**
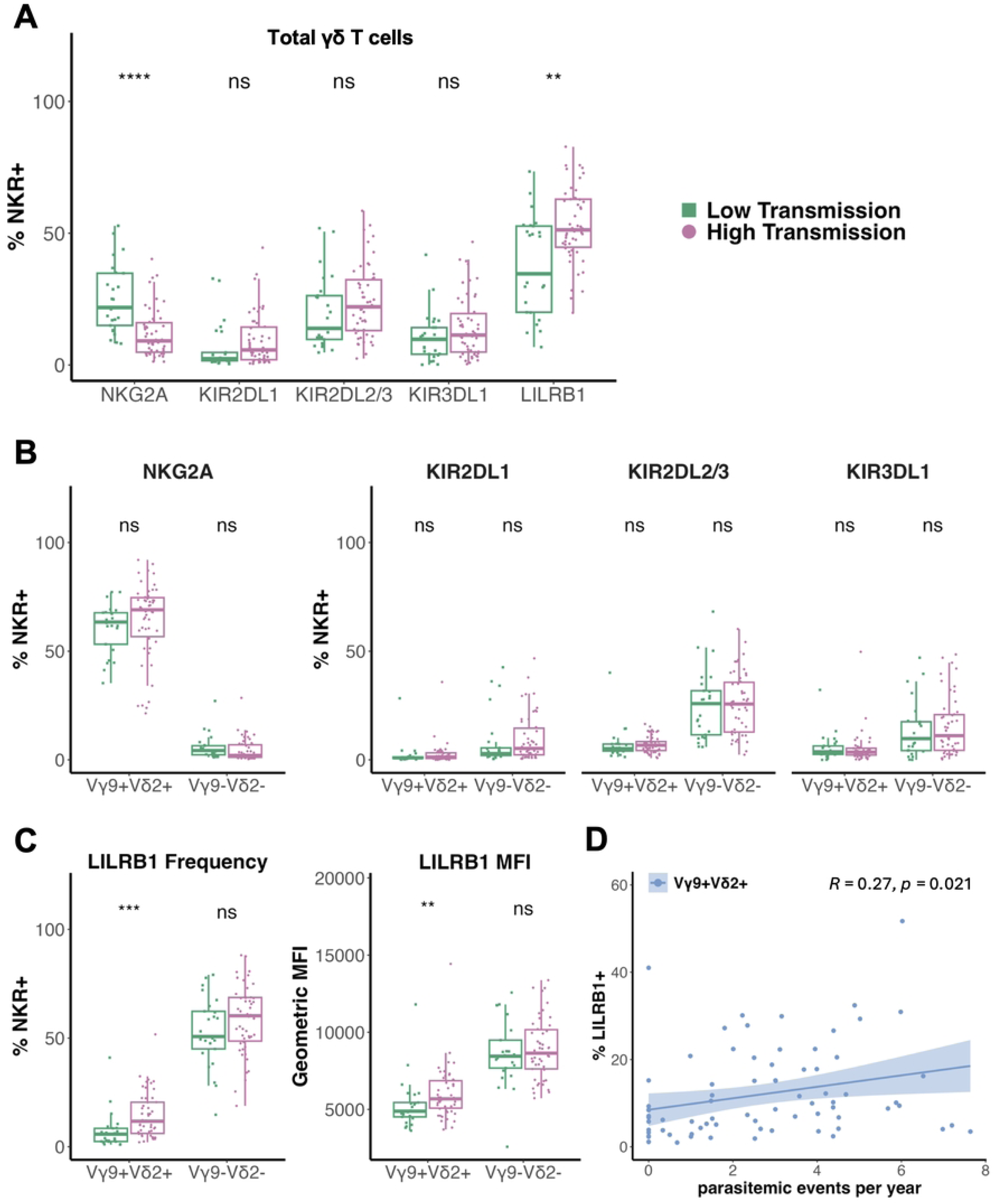
Malaria exposure has minimal impact on γδ T cell subset NKR expression. (**A**) *Ex vivo* frequency of inhibitory NK receptors on total γδ T cells, (**B**) frequency of NKG2A^+^ (left) and KIR^+^ (right) Vγ9^+^Vδ2^+^ and Vγ9^−^Vδ2^−^ T cells, and (**C**) frequency (left) and geometric mean fluorescence intensity (gMFI) (right) of LILRB1^+^ Vγ9^+^Vδ2^+^ and Vγ9^−^Vδ2^−^ T cells from individuals living in low and high malaria transmission sites. Study site groups were compared with an unpaired Wilcoxon rank sum test. For all analyses, a 2-tailed P value < 0.05 was considered significant (**: p <= 0.01, ***: p <= 0.001, ****: p <= 0.0001). (**D**) Frequency of LILRB1^+^ Vγ9^+^Vδ2^+^ T cells as a function of parasitemic events per year. Regression line is a standard linear model with the 95% confidence interval shaded in blue.

In contrast, LILRB1 was expressed more frequently on Vγ9^+^Vδ2^+^ T cells from highly malaria-exposed Tororo participants (median = 11.70%) than less malaria-exposed Jinja participants (median = 5.78%) (*p = 0.00024*, **Fig 3C**). Further, the cell surface density of LILRB1 was also higher in Tororo participants (median geometric mean fluorescence intensity (gMFI) = 5685) than Jinja participants (median gMFI = 4885) (*p = 0.009*, **Fig 3C**). Both the percentage of LILRB1-expressing cells and gMFI on Vγ9^+^Vδ2^+^ T cells correlated with the number of prior parasitemic events per year (**Fig 3D, S2 Fig**). These data echo similar observations in NK cells, in which LILRB1 expression is higher on CD56^neg^CD16^+^ NK cells—an atypical NK cell subset that expands in individuals with high malaria exposure—compared to CD56^dim^CD16^+^ NK cells [38]. Together, this suggests chronic malaria antigen exposure may drive an increase in LILRB1 expression; or alternatively, it may reflect preferential survival of LILRB1^+^ Vγ9^+^Vδ2^+^ T cells in the setting of parasitemia.

### NKG2A and KIR3DL1 expression associates with enhanced Vγ9^+^Vδ2^+^ T cell function

We next investigated the impact of inhibitory NK receptor expression on the antimalarial function of Vγ9^+^Vδ2^+^ T cells. We assessed both the TCR-mediated and CD16-mediated response of Vγ9^+^Vδ2^+^ T cells: we stimulated cells with (E)-4-Hydroxy-3-methyl-but-2-enyl pyrophosphate (HMBPP), a phosphoantigen produced by *Plasmodium* and a strong indirect ligand for the Vγ9^+^Vδ2^+^ TCR, as well as with plate-bound α-CD16 antibody which induces CD16 crosslinking. We measured degranulation by surface mobilization of CD107a as well as production of pro-inflammatory cytokines IFNγ and TNFα (**S3 Fig**). Further, we tested whether disrupting the binding of inhibitory NK receptors to their HLA class I (HLA-I) ligands with a pan-HLA-I blocking antibody alters the Vγ9^+^Vδ2^+^ T cell response to HMBPP.

First, we examined the impact of expression of NKG2A, the most prevalent inhibitory NK receptor on Vγ9^+^Vδ2^+^ T cells. In response to both HMBPP stimulation (**Fig 4A)** and CD16 crosslinking (**S4A Fig**), we found that Vγ9^+^Vδ2^+^ T cells expressing NKG2A degranulated and produced cytokines more frequently than NKG2A^−^ cells. Surprisingly, however, NKG2A^+^ Vγ9^+^Vδ2^+^ T cells produced IFNγ less frequently in response to HMBPP with HLA-I blockade than HMBPP alone (**Fig 4B**). NKG2A expression on Vδ2^+^ T cells has previously been shown to associate with more potent antitumor effector function in cancer models, lending to the idea that these cells are educated by the NKG2A-HLA interaction to recognize self and permit tumor killing, thereby becoming “licensed” in a manner similar to NK cells [39]. Our data indicate that expression of NKG2A may similarly license γδ T cells for more potent responsiveness to pathogens.

**Fig 4.**
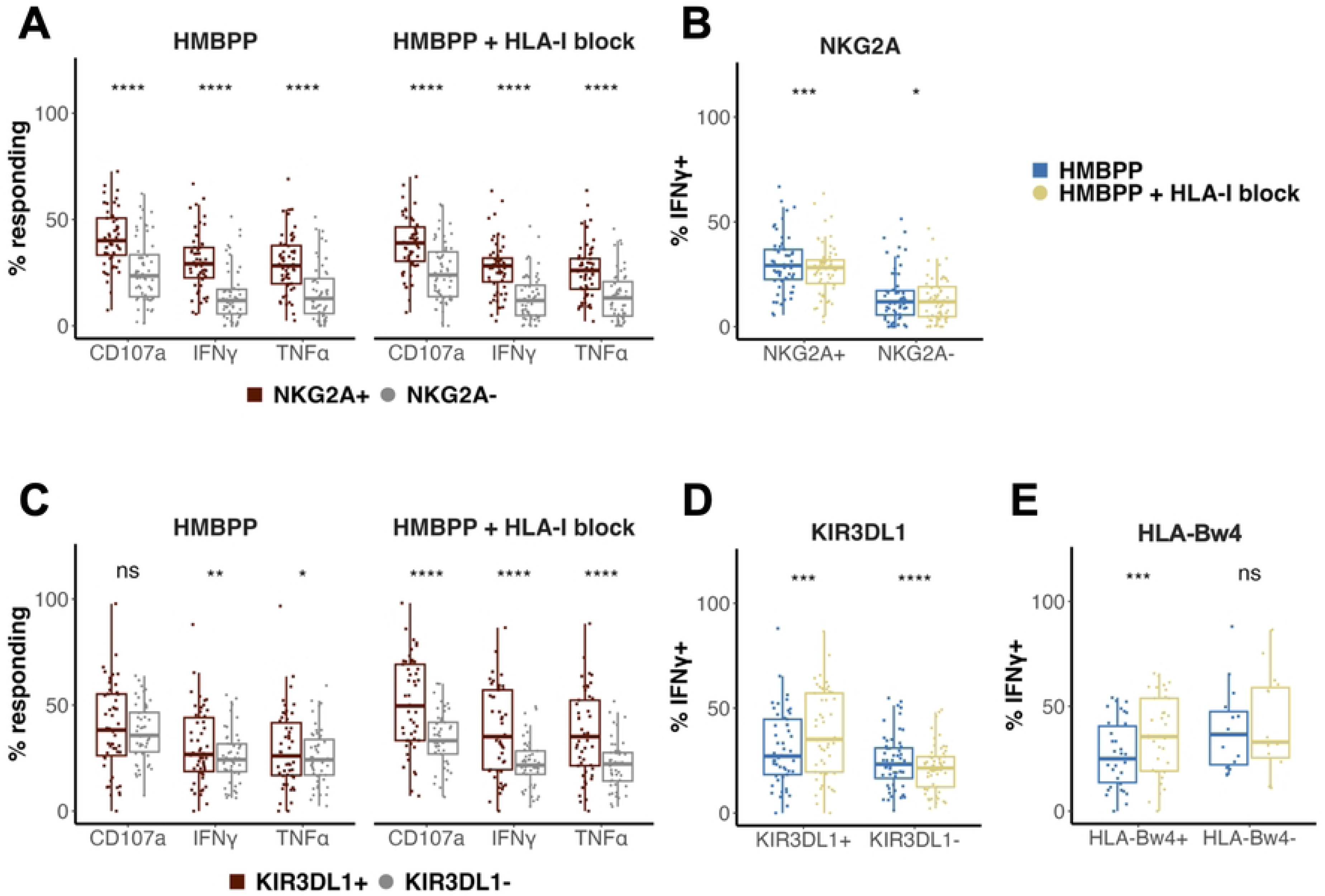
NKG2A and KIR3DL1 associate with potent cytotoxic and pro-inflammatory Vγ9^+^Vδ2^+^ T cell response. (**A**) Functional response of NKG2A**^+^** vs. NKG2A^−^ Vγ9**^+^**Vδ2**^+^** T cells, as measured by surface mobilization of CD107a and production of IFNγ and TNFα upon a 5-hour stimulation with malaria phosphoantigen alone (HMBPP, n = 56) or following a 30-minute pre-incubation with a class I HLA blocking antibody (HMBPP + HLA-I block, n = 56). (**B**) Frequency of IFNγ**^+^** NKG2A**^+^** or NKG2A^−^ Vγ9**^+^**Vδ2**^+^** T cells stimulated with HMBPP vs. HMBPP + HLA-I block (NKG2A**^+^**: n = 56; NKG2A^−^: n = 56). (**C**) Functional response of KIR3DL1**^+^** vs. KIR3DL1^−^ Vγ9**^+^**Vδ2**^+^** T cells stimulated with HMBPP (n = 50) or HMBPP + HLA-I block (n = 49). (**D**) Frequency of IFNγ**^+^** KIR3DL1**^+^** or KIR3DL1^−^ Vγ9**^+^**Vδ2**^+^** T cells stimulated with HMBPP vs. HMBPP + HLA-I block (KIR3DL1**^+^**: n = 49; KIR3DL1^−^: n = 56). (**E**) Frequency of IFNγ**^+^** KIR3DL1**^+^** Vγ9**^+^**Vδ2**^+^** T cells grouped by individuals with (n = 34) or without (n = 15) the HLA-Bw4 allele in HMBPP vs. HMBPP + HLA-I block conditions. All data were background-subtracted with matched unstimulated samples, and groups were compared with a paired Wilcoxon rank sum test. For all analyses, a 2-tailed P value < 0.05 was considered significant (*: p <= 0.05, **: p <= 0.01, ***: p <= 0.001, ****: p <= 0.0001). Data were included if parent gate contained a minimum of 10 cells.

While Vγ9^+^Vδ2^+^ T cells expressing KIR3DL1 were far less abundant than those expressing NKG2A, they exhibited a similar pattern, with KIR3DL1^+^ cells producing IFNγ and TNFα more frequently than KIR3DL1^−^ cells, although this difference was more subtle (**Fig 4C, S4A Fig**). The enhanced function of KIR3DL1-expressing cells was augmented by pre-incubation with a class I HLA blocking antibody (**Fig 4C**). This augmentation was specific to cells expressing KIR3DL1, and was not seen in KIR3DL1^−^ Vγ9^+^Vδ2^+^ T cells, suggesting a ligand-specific interaction (**Fig 4D**). KIR3DL1^−^ Vγ9^+^Vδ2^+^ T cells, in contrast, showed a reduced response upon HLA blockade—likely associated with NKG2A which is expressed on most KIR^−^ cells (**Fig 4B**). To further explore the specificity of the HLA blockade effect on KIR3DL1+ cells, we grouped participants according to whether they possessed or lacked the HLA-Bw4 ligand for KIR3DL1. We found that the enhanced responsiveness of KIR3DL1^+^ Vγ9^+^Vδ2^+^ T cells to stimulation upon HLA blockade was seen only in HLA-Bw4^+^ individuals (**Fig 4E**), whose γδ T cells were “educated” by HLA-Bw4. Together, these data suggest that the interaction of KIR3DL1 with its ligand, HLA-Bw4, may license cells to respond downstream to activating signals.

### KIR2DL1, KIR2DL2/3, and LILRB1 expression associates with attenuated Vγ9^+^Vδ2^+^ T cell function

In the case of the remaining two inhibitory KIR, we observed the opposite pattern. In response to HMBPP stimulation and CD16 crosslinking, Vγ9^+^Vδ2^+^ T cells expressing KIR2DL1 or KIR2DL2/3 degranulated and produced cytokines much less frequently than cells lacking these KIR (**Fig 5A**, **S4A Fig**). The functional response of KIR2DL1^+^ and KIR2DL2/3^+^ Vγ9^+^Vδ2^+^ T cells to HMBPP stimulation was minimally altered under HLA-I blockade.

**Fig 5.**
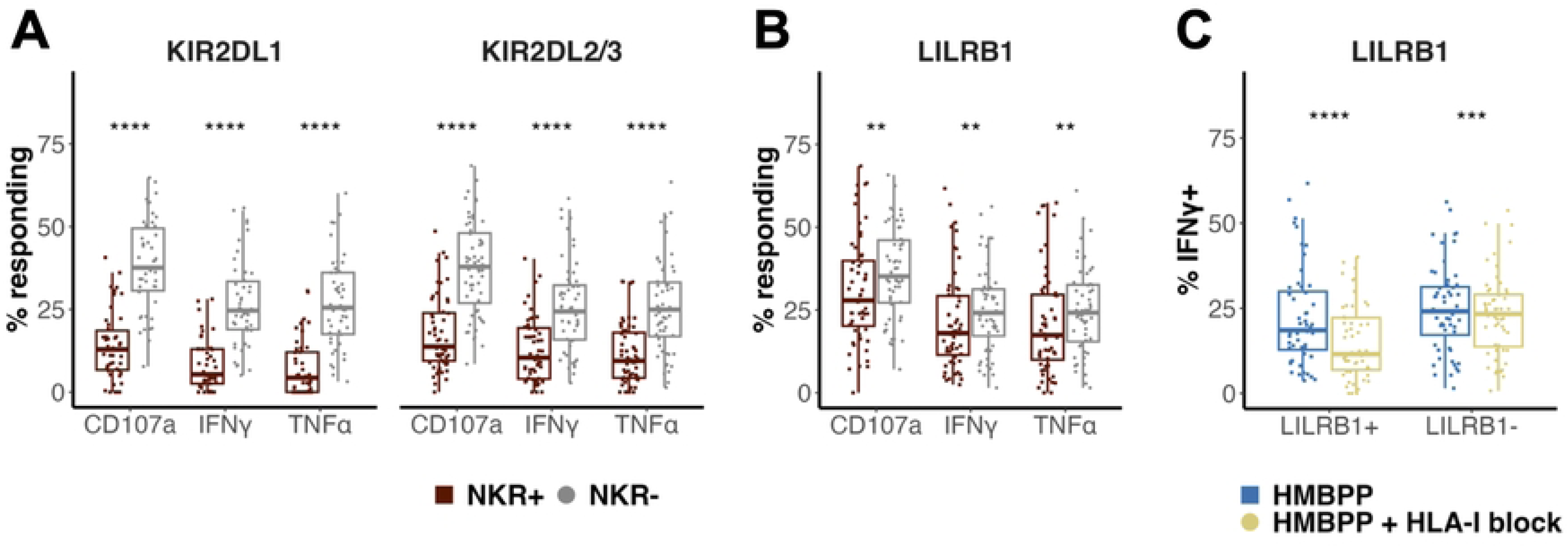
KIR2DL1, KIR2DL2/3, and LILRB1 associate with reduced Vγ9^+^Vδ2^+^ T cell response. (**A**) Functional response of KIR**^+^** vs. KIR^−^ Vγ9**^+^**Vδ2**^+^** T cells, as measured by surface mobilization of CD107a and production of IFNγ and TNFα upon a 5-hour stimulation with malaria phosphoantigen HMBPP (KIR2DL1, n = 46; KIR2DL2/3, n = 56). (**B**) Functional response of LILRB1**^+^** vs. LILRB1^−^ Vγ9**^+^**Vδ2**^+^** T cells stimulated with HMBPP (n = 56). (**C**) Frequency of IFNγ**^+^** LILRB1**^+^** or LILRB1^−^ Vγ9**^+^**Vδ2**^+^** T cells stimulated with HMBPP vs. HMBPP + HLA-I block (LILRB1**^+^**: n = 52; LILRB1^−^: n = 56). All data were background-subtracted with matched unstimulated samples, and groups were compared with a paired Wilcoxon rank sum test. For all analyses, a 2-tailed P value < 0.05 was considered significant (**: p <= 0.01, ***: p <= 0.001, ****: p <= 0.0001). Data were included if parent gate contained a minimum of 10 cells.

Lastly, we investigated the functional response of Vγ9^+^Vδ2^+^ T cells expressing LILRB1, which can notably bind all HLA-I types [40]. We observed that LILRB1^+^ Vγ9^+^Vδ2^+^ T cells degranulated less frequently in response to both stimuli and produced cytokines less frequently in response to HMBPP stimulation compared to cells lacking LILRB1 (**Fig 5B, S4A Fig**). Surprisingly, however, this inhibitory effect was augmented by HLA-I blockade, with LILRB1^+^ Vγ9^+^Vδ2^+^ T cells producing IFNγ less frequently under blockade (**Fig 5C**). To a lesser extent, LILRB1^−^ Vγ9^+^Vδ2^+^ T cells also responded less under HLA-I blockade, likely due to NKG2A as described above. In all, these results suggest that LILRB1, KIR2DL1, and KIR2DL2/3 are each associated with inhibited Vγ9^+^Vδ2^+^ T cell functional responses, and that disrupting binding to their HLA-I ligands is not sufficient to rescue their activity.

## DISCUSSION

Here, we found that γδ T cells frequently express inhibitory NK receptors, but their expression differs starkly between subsets. Vγ9^+^Vδ2^+^ T cells more frequently expressed NKG2A while Vδ2^−^ and Vγ9^−^Vδ2^+^ T cells predominantly expressed LILRB1 or a combination of KIRs, pointing to divergent mechanisms of inhibitory control. While malaria exposure intensity influenced the relative frequency of γδ T cell subsets in peripheral blood, it did not influence NKR expression within a given subset, with the exception of LILRB1 expression by Vγ9^+^Vδ2^+^ T cells. We further showed that expression of inhibitory NK receptors on Vγ9^+^Vδ2^+^ T cells is associated with differences in responsiveness to malaria antigen stimulation. Vγ9^+^Vδ2^+^ T cells expressing NKG2A and to a lesser extent KIR3DL1 had a more robust response to both TCR-mediated activation by phosphoantigens and to direct CD16 ligation and crosslinking. In contrast, the responsiveness of Vγ9^+^Vδ2^+^ T cells expressing KIR2DL1, KIR2DL2/3 and LILRB1 was attenuated. These findings support the hypothesis that inhibitory NK receptors are involved in regulating γδ T cell effector responses.

Our most striking finding is that NKG2A and KIR3DL1 expression is associated with a more potent Vγ9^+^Vδ2^+^ T cell response to TCR and CD16 stimulation. It has been proposed that NKG2A on Vγ9^+^Vδ2^+^ T cells might follow the “licensing” or “arming” model of NK cells, in which inhibitory receptors are required to educate the cell to recognize self before it is able to respond beyond a certain threshold of activation [39,41,42]. There is evidence of this in cancer, as NKG2A^+^ Vγ9^+^Vδ2^+^ T cells exhibit hyper-responsiveness to tumor cells compared to NKG2A^−^ cells [39]. To our knowledge, there is no such evidence for KIR3DL1 licensing Vγ9^+^Vδ2^+^ T cells, yet we found that KIR3DL1-expressing Vγ9^+^Vδ2^+^ T cells produce more IFNγ and TNFα in response to stimulation. Disrupting HLA binding in HLA-Bw4^+^ individuals further enhanced both degranulation and cytokine production in response to HMBPP stimulation, which points to education of KIR3DL1^+^ cells by its HLA ligand that enables a strong response in the absence of binding. Therefore, our study supports the currently limited evidence for γδ T cell education mediated by inhibitory receptors and expands its biological context beyond antitumor effector function to encompass responding to pathogens.

In contrast, we found that KIR2DL1^+^, KIR2DL2/3^+^, and LILRB1^+^ Vγ9^+^Vδ2^+^ T cells were significantly less responsive to stimulation by malaria phosphoantigens and by direct ligation of CD16. Although these populations constitute a minority of this γδ T cell subset, this finding nonetheless suggests a model whereby expression of these inhibitory receptors dampens Vγ9^+^Vδ2^+^ T cell responses and lessens their parasite-killing capacity. Supporting this model, we previously showed in a large immunogenetic study that possession of KIR2DL1 (the most strongly inhibitory KIR) along with its ligand HLA-C2 was associated with a higher risk of *P. falciparum* parasitemia [22]. Furthermore, a separate study in a Thai population revealed a risk associating KIR2DL3 and HLA-C1 with severe cerebral malaria, an inflammatory complication of the disease [43]. Intriguingly, *P. falciparum* itself encodes a family of proteins called RIFINs that are expressed on the surface of infected erythrocytes and can directly ligate inhibitory receptors including KIR2DL1, LILRB1, and LAIR1 [44]. These inhibitory interactions with RIFINs suppress immune cell activation, enabling the parasite to escape killing, and LILRB1-binding parasite strains have been associated with more severe malaria [23]. Remarkably, naturally occurring antibodies containing RIFIN-binding domains of LILRB1 have been identified in individuals living in malaria-endemic regions [45–48]. These antibodies are able to block the LILRB1-RIFIN interaction and permit the host immune response, demonstrating the clear evolutionary importance of the LILRB1 signaling pathway in malaria immunopathogenesis [45–48]. KIR2DL1 was recently implicated as another RIFIN-binding receptor that can suppress killing by NK cells (and perhaps γδ T cells as well), enabling parasite persistence [49]. These inhibitory receptors appear to lie at the crux of parasite immune evasion and host immune control, and present an exciting avenue for future work to determine their therapeutic potential.

With the exception of LILRB1 on Vγ9^+^Vδ2^+^ T cells, the intensity of prior malaria exposure was not associated with the frequency of inhibitory NK receptor expression within a given γδ T cell subset. Rather, γδ T cell subsets exhibit distinct patterns of NK receptor expression, and malaria exposure results in overall shifts in the proportions of these subsets in peripheral blood. We found that a majority of Vγ9^+^Vδ2^+^ T cells express NKG2A, whereas both Vγ9^−^Vδ2^+^ and Vδ2^−^ T cells have markedly higher LILRB1 and KIR expression. NKG2A and KIR represent two structurally distinct but functionally complementary “schools” of inhibitory HLA class I receptors that educate NK cells and influence their functional potency [50]. NKG2A is an older and more conserved receptor, whereas KIR are more recently evolved and highly diversified [51]. Hence, our observation that Vγ9^+^Vδ2^+^ T cells exhibit divergent inhibitory NK receptor expression from the Vδ2^−^ and Vγ9^−^Vδ2^+^ subsets is congruent with the emerging view of Vγ9^+^Vδ2^+^ T cells as more innate-like, while Vδ1^+^ and Vγ9^−^Vδ2^+^ T cells more often exhibit adaptive-like effector profiles [14,16,32,33].

The role of inhibitory NK receptors in regulating Vγ9^−^Vδ2^−^ T cells remains ambiguous. They are the most prevalent circulating γδ T cell subset in highly malaria-exposed individuals, and clonally expanded cytotoxic Vδ1^+^ T cells are present at increased frequencies in malaria-exposed individuals in Mali [30]. A similar pattern has been observed in the context of controlled *Mycobacterium tuberculosis* (*Mtb*) infection; chronically *Mtb*-infected individuals have a higher proportion of CD8^+^ γδ T cells that primarily utilize the Vδ1 chain and express high levels of inhibitory KIRs as well as cytotoxic and cytolytic molecules [52]. It is thought that these cells represent a functional NK-like γδ T cell repertoire that stems from constant antigen exposure, which could play a role in other chronic infections like malaria. Meanwhile, Vγ9^+^Vδ2^+^ T cells expressing NKG2A could represent a reserve of ready-made effector cells as they maintain hyper-responsiveness compared to NKG2A^−^ cells despite their reduced frequencies in highly malaria-exposed individuals. Future mechanistic studies are needed to elaborate the parasite-killing function of these NK receptor-expressing γδ T cells.

Our study has several limitations. While we observed differences in the functional responses of Vγ9^+^Vδ2^+^ T cells that express or lack a given NK receptor, we cannot conclude that these differences are causally linked to that particular receptor. Other co-expressed NK receptors (including those not evaluated here), or associated transcriptional programming, may also contribute to modulation of the γδ T cell response. The means by which activating and inhibitory signals are integrated at the immunological synapse remains ambiguous, particularly for γδ T cells, which respond to a broad array of activating signals. While we showed in some cases that HLA-I blockade alters the response to stimulation, this blockade could disrupt numerous interactions with the many activating and inhibitory receptors that bind HLA-I. Future investigations using targeted blockade with NKR-specific antibodies could help to further dissect the impact of ligation of individual inhibitory receptors. Finally, in addition to TCR- and FcR-mediated activation, γδ T cells can receive activating signals through cytokine receptors, so it is possible that our functional assays capture bystander effects of cytokines produced by other cells in culture. However, this is likely mitigated by the brief duration of stimulation in our assay. Moreover, we have shown in a prior study that purified γδ T cells and those in whole PBMC respond in a comparable manner to CD16 stimulation [19].

In summary, we show that subsets of γδ T cells exhibit different patterns of inhibitory NK receptor expression *ex vivo*, and that expression of these NK receptors is associated with an altered response to stimulation. On Vγ9^+^Vδ2^+^ T cells, which are known to be highly functional in the response to primary malaria infection but contract and become less responsive in chronic repeated infection, we illustrate that expression of NKG2A and to a lesser extent KIR3DL1 licenses strong effector programs while KIR2DL1, KIR2DL2/3, and LILRB1 expression inhibits degranulation and cytokine production. While further work will be needed to understand the mechanism by which these receptors translate stimuli and modulate downstream functional outputs, these findings expand our knowledge of conventional inhibitory NK receptors expressed on γδ T cells and their role in regulating the immune response to malaria.

## METHODS

### STUDY SITES AND DESIGN

Details of the Program for Resistance, Immunology, Surveillance, and Modeling of Malaria in Uganda (PRISM Cohort Study) have been described elsewhere [26]. Briefly, children between 6 months and 10 years of age from 100 randomly selected households across three study sites in Uganda were enrolled and followed longitudinally. Participants received care at dedicated study clinics where blood was drawn and parasitemia was routinely monitored via thick blood smear every three months in addition to intervening symptomatic visits. The described experiments include samples from individuals from Walakuba (Jinja District; n = 25), a peri-urban region with an annual entomologic inoculation rate (aEIR) of 2.8, and individuals from Nagongera (Tororo District; n = 50), a rural region with an aEIR of 310 [26]. All PBMC samples included in this study were collected before the initiation of indoor residual spraying (IRS) of carbamate bendiocarb in Tororo in December 2014 [53].

Additionally, samples were selected such that each study site had mixed representation by individuals who were either homozygous or heterozygous for the HLA-C1 or HLA-C2 allotypes, and either possessed or did not possess the HLA-Bw4 allele (C1/C1, n = 30; C1/C2, n = 15; C2/C2, n = 30; Bw4^+^, n = 50; Bw4^−^, n = 25). The prevalence of KIR genotypes in the selected participants were as follows: 100% 2DL1, 58.7% 2DL2, 92% 2DL3, 98.7% 3DL1, 18.7% 2DS1, 52% 2DS2, 21.3% 2DS3, 12% 3DS1. CMV serostatus of all participants included in this study was determined using the Diamedix™ Immunosimplicity™ human anti-cytomegalovirus IgG ELISA kit (Fisher Scientific). All but one participant (participant ID: 3012) were CMV seropositive. All participants were HIV negative.

### ETHICAL APPROVAL

Written informed consent was obtained from all study participants or the guardian of participants under 18 years of age. Study protocols were approved by the Uganda National Council of Science and Technology, the Makerere University School of Medicine Research and Ethics Committee, and the University of California, San Francisco (UCSF) Committee on Human Research.

### SAMPLE PROCESSING

At regular quarterly visits, 3 to 10 ml of blood was collected with acid citrate dextrose tubes. Plasma was removed and frozen at –80 °C, and peripheral blood mononuclear cells (PBMC) were isolated via density gradient centrifugation (Ficoll-Histopaque; GE Life Sciences) and cryopreserved in liquid nitrogen before shipping to UCSF for downstream analysis.

### SURFACE AND INTRACELLULAR CYTOKINE STAINING

Frozen PBMC were thawed in RPMI 1640 (Gibco) supplemented with 10% v/v heat-inactivated FBS (GeminiBio), Pen/Strep (100 IU/ml Penicillin:0.01 mg/ml Streptomycin, GeminiBio), L-glutamine (4 mM, Gibco), HEPES (10 mM, Gibco), and DNAse (1.25 units/ml, Roche) and counted, each sample passing a viability threshold of >90%. Cells were divided such that a minimum of 750,000 cells per sample were subjected either to immediate *ex vivo* surface staining or overnight rest at 2M cells/ml at 37 °C before stimulation followed by surface and intracellular cytokine staining. SpectraComp^Ⓡ^ compensation beads (Slingshot Bio) were used for all single stain controls except for viability and NKG2A, which used PBMC (viability used a 50% mixture of live and heat-killed cells). For the surface stain, cells were washed twice with PBS in a V-bottom 96-well plate, stained for 30 min at room temperature with the antibodies listed in **S1 Table** diluted in FACS staining buffer (R&D Systems), then washed 3 times with PBS. For intracellular cytokine staining, cells were permeabilized with the BD Cytofix/Cytoperm^TM^ fixation/permeabilization solution at 4 °C for 20 min, washed 2 times with BD Perm/Wash^TM^ buffer, stained with the intracellular stain antibodies listed in **S1 Table** for 1 hour at 4 °C, then washed 3 more times with BD Perm/Wash^TM^ buffer. All stained cells were fixed with 1% PFA in PBS and stored at 4 °C until acquisition. Fluorescence minus one (FMO) controls were generated for KIR2DL1, KIR2DL2/3, KIR3DL1, and LILRB1. PBMC from a single adult naive control donor were fully stained and used as a technical control for multiple processing batches.

### HMBPP STIMULATION, HLA-I BLOCK, CD16 STIMULATION

Using a master mix in serum-free RPMI for all stimulation conditions, cells were stained with an anti-CD107a antibody and subjected to protein transport inhibition using 1:1500 v/v Brefeldin A (BD GolgiPlug™, undiluted) and 1:1500 v/v Monensin (BD GolgiStop™, undiluted) at the time of stimulation. For HMBPP stimulation, cells were stimulated with 15 nM (E)-1-Hydroxy-2-methyl-2-butenyl 4-pyrophosphate (HMBPP) (Sigma-Aldrich) in a U-bottom 96-well plate for 5 hours at 37 °C and 5% CO_2_. For samples receiving HLA block, a pan-HLA-I-specific mouse IgG1 anti-human antibody (DX17 [54,55], BD) was added at a concentration of 0.01 mg/ml in serum-free media to each sample and pre-incubated for 30 min at 37 °C before washing 3 times with PBS and adding the HMBPP stimulation cocktail. A corresponding unstimulated condition was included for every sample, using all the above listed reagents except HMBPP and the HLA blocking antibody. For CD16 crosslinking, uncoated Nunc™ MaxiSorp™ ELISA plates (Biolegend) were washed twice with PBS and coated overnight at 4 °C with either anti-CD16 antibody (3G8, Biolegend) or an isotype control IgG1 antibody (MOPC-21, Biolegend) at a concentration of 6 μg/ml in 0.1M carbonate buffer (pH 9.6). The coated plate was then washed twice with PBS and blocked with RPMI complete medium for 10 min at room temperature. Blocking media was replaced with stimulation mastermix and cells were added for a stimulation of 5 hours at 37 °C and 5% CO_2_. After all 5 hour stimulations, samples were stored overnight at 4 °C before subsequent staining.

### SPECTRAL FLOW CYTOMETRY ANALYSIS

Populations of interest were gated on single cells, Zombie UV negative, CD19^−^, CD14^−^ lymphocytes that were CD3^+^ and pan-γδTCR^+^ for γδ T cells (**S1 Fig, S3 Fig**). At least 300,000 events per sample were collected for *ex vivo* samples and at least 100,000 events were collected for stimulated samples. Samples were analyzed on a 5-laser Aurora spectral flow cytometer (Cytek^Ⓡ^) with the SpectroFlo^Ⓡ^ (v3.1.0) software (Cytek^Ⓡ^). Spectral data was unmixed in SpectroFlo^Ⓡ^ and analyzed using FlowJo (v10.10.0) (BD).

### STATISTICAL METHODS

All statistical analyses were performed using R (v4.4.1) and RStudio (v2024.12.0+467), utilizing the dplyr (v1.1.4) and ggplot2 (v3.5.2) packages. Unless otherwise specified, phenotypic comparisons were performed using an unpaired Wilcoxon rank sum test, and functional comparisons were performed using a paired Wilcoxon rank sum test. Box plots show medians with Tukey whiskers. For all analyses, a 2-tailed P value < 0.05 was considered significant (*: p <= 0.05, **: p <= 0.01, ***: p <= 0.001, ****: p <= 0.0001). Background subtraction was applied to all samples stimulated with HMBPP or HMBPP + HLA-I block, subtracting frequencies from specimen-matched unstimulated samples. Background subtraction was not applied to CD16-stimulated samples as limited cell numbers prevented complete specimen matching of isotype control samples. CD16-stimulated and isotype control samples were adjusted using *ex vivo* CD16 expression frequencies such that reported functional responses represented the percentage of cells capable of responding to CD16 crosslinking. Briefly, the frequencies of responding cells were divided by the decimal percentage of CD16^+^ cells in the corresponding *ex vivo* parent gate.

### HIGH DIMENSIONAL ANALYSIS OF FLOW CYTOMETRY DATA

Dimensionality reduction was performed using R (v4.4.1) and RStudio (v2024.12.0+467), utilizing the CytoNorm (v2.0.9) and Cyclone (v0.1.0) packages [36,56]. γδTCR^+^ cells only were extracted as ungated FCS files using FlowJo (v10.10.0) (BD). Only relevant flow channels for markers downstream of pan-γδTCR were used for clustering, normalization and UMAP generation. FCS files were read into R as a flowFrame using the flowCore (v2.18.0) package. The CytoNorm model was trained with the adult naive control sample used as a technical control for all processing weeks (n = 4), clustering a subsample of 6000 cells using xdim = 15, ydim = 15, and clusters = 10 (FlowSOM (v2.14.0)), and computing 99 quantiles within each cluster to normalize to the goal quantile (average). This learned normalization model was then applied to the remaining samples, outputting adjusted FCS files for use in the Cyclone pipeline.

Using Cyclone, normalized data (excluding controls) were arcsinh transformed using a cofactor of 6000. UMAP was calculated using Nearest Neighbors = 30, minimum distance = 0.1, spread = 0.1, and learning rate = 0.5. Finally, Cyclone outputs were compiled into an S4 object using the SingleCellExperiment (v1.28.1) package and feature maps were generated with dittoSeq (1.18.0).

## SUPPORTING INFORMATION

**S1 Table. Antibody details.**

**S1 Fig. Gating schematic for NKR phenotyping of γδ T cells.** Lymphocytes were gated first, followed by single cells, live cells, and then γδ T cells. γδ T cells were gated on CD3^+^CD14^−^ CD19^−^ cells, followed by pan-γδTCR^+^, then divided into subsets based on expression of Vγ9 and Vδ2 TCR chains. Representative gating of NKRs on Vγ9^+^Vδ2^+^ T cells. KIRs and LILRB1 were gated with FMO samples. Identical gates were applied to all *ex vivo* phenotyping samples.

**S2 Fig. Surface density of LILRB1 on Vγ9^+^Vδ2^+^ T cells increases with parasitemic events.**

Geometric MFI of LILRB1 on Vγ9^+^Vδ2^+^ T cells as a function of parasitemic events per year. Regression line is a standard linear model with the 95% confidence interval shaded in blue.

**S3 Fig. Gating schematic for functional assays.** Lymphocytes were gated first, followed by single cells, live cells, and then γδ T cells. γδ T cells were gated on CD3^+^CD14^−^CD19^−^ cells, followed by pan-γδTCR^+^, then divided into subsets based on expression of Vγ9 and Vδ2 TCR chains. Representative gating of response markers on Vγ9^+^Vδ2^+^ T cells. CD107a, IFNγ, and TNFα were gated with corresponding unstimulated/isotype control samples. Identical gates were applied to all HMBPP-, HMBPP + HLA-I block-, and CD16-stimulated samples.

**S4 Fig. NKR expression affects Vγ9^+^Vδ2^+^ T cell response to CD16 crosslinking.** (**A**) Functional response of NKR^+^ vs. NKR^−^ Vγ9^+^Vδ2^+^ T cells, as measured by surface mobilization of CD107a and production of IFNγ and TNFα upon a 5-hour stimulation with plate-bound α-CD16 crosslinking antibody. Data is not background subtracted. NKG2A: n = 69; KIR3DL1: n = 51; KIR2DL1: n = 50; KIR2DL2/3: n = 68; LILRB1: n = 67. (**B**) Background response of NKR^+^ vs. NKR^−^ Vγ9^+^Vδ2^+^ T cells stimulated with plate-bound isotype control IgG1 antibody. NKG2A: n = 45; KIR3DL1: n = 31; KIR2DL1: n = 32; KIR2DL2/3: n = 44; LILRB1: n = 43. NKR expression categories were compared via paired Wilcoxon rank sum test, with datapoints paired by donor. For all analyses, a 2-tailed P value < 0.05 was considered significant (*: p <= 0.05, **: p <= 0.01, ***: p <= 0.001, ****: p <= 0.0001). Data were included if parent gate contained a minimum of 10 cells.

**S1 Data. Values used to construct all graphs in the manuscript are listed in tables, labeled with their corresponding figure number and panel.**

## ACKNOWLEDGEMENTS

We are incredibly grateful to all study participants and their families for making this work possible. We would like to thank all the study team members for their invaluable contributions. Additionally, we thank Victoria Tran, Kyla Foster, Dr. Shayan Avanessian, Dr. Sara Suliman and Dr. Oscar Aguilar for their insights and guidance while preparing this manuscript. We would also like to acknowledge the Core Immunology Lab for their assistance with flow cytometry equipment.

